# CParty: Hierarchically Constrained Partition Function of RNA Pseudoknots

**DOI:** 10.1101/2023.05.16.541023

**Authors:** Luke Trinity, Mateo Gray, Sebastian Will, Yann Ponty, Ulrike Stege, Hosna Jabbari

## Abstract

Biologically relevant RNA secondary structures are routinely predicted by efficient dynamic programming algorithms that minimize their free energy. Starting from such algorithms, one can devise partition function algorithms, which enable stochastic perspectives on RNA structure ensembles. As most prominent example McCaskill’s partition function algorithm is derived from pseudoknot-free energy minimization. While this algorithm became hugely successful for the stochastic analysis of pseudoknot-free RNA structure, as of yet there exists only one pseudoknotted partition function implementation, which covers only simple pseudoknots and comes with a borderline-prohibitive complexity of *O*(*n*^5^) in the RNA length *n*. In this article, we develop a partition function algorithm corresponding to the hierarchical pseudoknot prediction of HFold, which performs exact optimization in a realistic pseudoknot energy model. In consequence, our algorithm CParty carries over HFold’s advantages over classical pseudoknot prediction to stochastic analysis. In only cubic time, it computes the hierarchically constrained partition function over pseudoknotted density-2 structures *G* ∪ *G*^*′*^, composed of pseudoknot-free parts *G* and *G*^*′*^, where *G* is given. Thus, it follows the common hypothesis of hierarchical pseudoknot formation, where pseudoknots form as tertiary contacts only after a first pseudoknot-free ‘core’ *G*. Like HFold, CParty is very efficient, achieving the low complexity of the pseudoknot-free algorithm. Finally, by computing pseudoknotted ensemble energies, we unveil kinetics features of a therapeutic target in SARS-CoV-2.

**Availibility:** CParty is available at https://github.com/HosnaJabbari/CParty.

## Introduction

RNA molecules play a vital role in cellular processes; many possess functional structures [12, 25, 32, 47, 48]. As experimental methods to detect RNA structure are time consuming and costly, computational methods for predicting RNA structure have become indispensable. We focus on the accurate prediction of RNA secondary structure (2D), which in turn sheds light on the 3D structure of the RNA. Various algorithms have been developed to tackle this problem, aiming to predict the most energetically favorable structure based on thermodynamic models and empirical data [7, 36, 37, 38, 39, 52]. Two widely used thermodynamics-based approaches for predicting RNA secondary structure are by Zuker [52] and McCaskill [31].

The Zuker algorithm finds the minimum free energy (MFE) structure among all possible pseudoknot-free structures for the given RNA sequence [52]. RNA secondary structure prediction is NP-hard [1, 27] and even inapproximable [42] when pseudoknots are allowed. Existing efficient algorithms for exact prediction of pseudoknotted RNA secondary structure handle only restricted classes of structures, trading off run-time and structure complexity [10, 23, 38].

Offering a stochastic perspective on the entire ensemble of possible pseudoknot-free structures of an RNA, the McCaskill algorithm computes *partition functions*. Algorithmically it has strong parallels to the pseudoknot-free MFE algorithm by Zuker, since both algorithms decompose the same structure space in their dynamic programming scheme. Generally, there is a one-to-one correspondence between the search spaces considered by partition function algorithms, such as McCaskill’s [31], and MFE algorithms, provided they are unambiguous and complete [33]. This correspondence also extends to pseudoknotted structure spaces. Consequently, the run-time vs. structure complexity trade-offs that were discussed for pseudoknot MFE algorithms like [10, 23, 38] are mirrored in (hypothetical) corresponding partition function algorithms. So far, the only pseudoknot partition function algorithm, which realizes this idea, is due to Dirks and Pierce [15]. Their algorithm (D&P) handles the restricted class of simple pseudoknots. While it is implemented in NUPACK, its practical application is limited by the algorithm’s *O*(*n*^5^) time and *O*(*n*^4^) space complexity.

To address the high time and space complexity of other pseudoknot prediction algorithms, we previously developed HFold [21, 22]. HFold efficiently predicts potentially pseudoknotted RNA secondary structures that extend a given pseudoknot-free secondary structure, accurately predicting exact optima within a full RNA energy model. By following this principle, HFold becomes the first MFE algorithm that adheres to the *hierarchical folding hypothesis*. This hypothesis suggests that RNA initially folds into a pseudoknot-free structure, and then additional bases pair to further lower the MFE of the structure, possibly forming pseudoknots [43]. Hierarchical folding has been experimentally observed in the formation of pseudoknotted structures [11], including frameshift stimulating pseudoknots [8].

HFold has only cubic time and quadratic space complexity. This means it is as efficient as pseudoknot-free prediction algorithms and, for example, is faster than CCJ [9] or D&P [15] by a quadratic factor. HFold takes a pseudoknot-free structure *G* as input, and predicts a pseudoknot-free structure *G*^′^ such that *G* ∪ *G*^′^ has minimum free energy among all *density-2* structures [22] (Figure 1; see also Definitions). The class of density-2 structures allows for arbitrary depth and length of nested pseudoknots including H-type pseudoknots and kissing hairpins. This class encompasses structures not handled by CCJ and is more comprehensive than the structure class described by the partition function algorithm by Dirks and Pierce [15]. Since the selection of the non-pseudoknotted partial structure *G* is crucial in hierarchical folding, previous work [20, 45] has identified promising techniques for selecting *G*. These techniques involve computing energetically favorable pseudoknot-free structures or choosing partial structures compatible with chemical modification data, such as SHAPE reactivity.

**Fig. 1.**
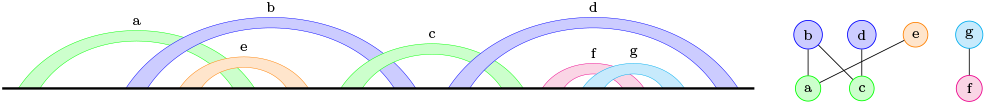
Bands of a bisecondary structure (left) and their crossing graph (right). Each band represents a set of pairwise nested base pairs; all base pairs within one band cross the exact same other base pairs or bands in the structure. Removing the orange band leaves a density-2 configuration of bands: no two bands within any of the two connected components are nested.

The main objective of this work is to develop and study a partition function counterpart to HFold. To achieve this, we present CParty, a *constrained partition function* (CPF) algorithm that considers possibly pseudoknotted density-2 structures.

The primary challenge in constructing the CParty algorithm is that HFold decomposes density-2 structures in a non-trivially redundant manner. To address this, we have resolved all ambiguities in the decomposition process. This careful preparation step, finally enables deriving the CParty algorithm by a systematic exchange of algebras (cf. ADP [17]).

Similar to HFold, CParty takes an RNA sequence and an input structure *G*. Then, it calculates the hierarchically constrained partition function over *G* ∪ *G*^′^, where *G*^′^ is pseudoknot-free and *G* ∪ *G*^′^ is density-2.

To clarify, CParty is designed to compute hierarchically constrained partition functions of RNAs, which ensures very high efficiency. In some applications, this hierarchical approach can be advantageous, as it allows for the integration of prior knowledge. Additionally, its efficiency makes it suitable for iterated use in meta-strategies (cf. [20]).

### Contributions

We introduce the novel hierarchical constraint partition function algorithm CParty as a counterpart to HFold. CParty decomposes the density-2 structure class completely and, in contrast to HFold, unambiguously. We implement CParty to perform realistic computations using a full-featured pseudoknot energy model (HotKnots 2.0 [4]), thoroughly scrutinize the implementation, and study its properties. Through empirical time complexity analysis, we demonstrate that CParty outperforms the only other existing pseudoknotted partition function algorithm in NUPACK. Applying our novel tool to the SARS-CoV-2 frameshift element, we compute constrained ensemble energies and unveil a key kinetic transition of its pseudoknot [24].

## Methods

### RNA Secondary Structure

An RNA *sequence* of length *n*, known as the *primary structure* of RNA, is represented as a string in {*A, C, G, U*}^*n*^. Its *secondary structure* is a set of base pairs *i*.*j*, where 1 ≤ *i < j* ≤ *n*, and each *base* 1 ≤ *i* ≤ *n* occurs in at most one pair (no triplets). A secondary structure is called *crossing* or *pseudoknotted* if there are at least two base pairs, *i*.*j* and *i*^′^.*j*^′^, that *cross* each other (i.e. *i < i*^′^ *< j < j*^′^ or *i*^′^ *< i < j*^′^ *< j*). Otherwise, it is called *pseudoknot-free* or *non-crossing*. The base pairs *i*.*j* and *i*^′^.*j*^′^ are *nested* if *i < i*^′^ *< j*^′^ *< j* or *i*^′^ *< i < j < j*^′^. Given a secondary structure *G* that pairs *i, bp*_*G*_(*i*) denotes the other end of the base pair of *i* in *G*; similarily, *bp*(*i*) refers to the other end in *G* ∪ *G*^′^.

We briefly review additional features of non-crossing structures. Due to space restrictions, we refer to the literature for details [15, 22]. Pseudoknotted structures contain subsets of base pairs called *bands* [15]. Bands are maximal subsets of base pairs that are pairwise nested, with each base pair crossing exactly the same base pairs in the remaining structure.

The literature distinguishes various classes of crossing structures that encompass different band crossing patterns, such as simple pseudoknots, kissing hairpins, *k*-knots, and genus *g*. In this work, we focus on a specific subclass of *bisecondary* structures [16, 18, 49], which can be decomposed into two pseudoknot-free secondary structures. Specifically, we consider the subclass of density-2 structures defined by Jabbari et al. [22] to precisely describe the search space of HFold. We illustrate a density-2 structure in Fig. 1. For a more formal intuition, envision the *crossing graph* representing the bands (nodes) and their crossing relations (edges). Bisecondary structures correspond to bipartite graphs; in density-2 structures, no two bands of the same connected component can be nested.

We require additional technical definitions from HFold’s description [22]: In density-2 structures, a *region* [*i, j*] (denoting positions *i, i* + 1, …, *j* is *closed*, either if *i* and *j* are paired, or if they are transitively connected due to a chain of crossing bands. In the latter case, *i* and *j* are the left and right ends of a *pseudoloop*, which is closed by base pairs of *i* and *j* as well as the outer base pairs of the other bands in the chain. For example, in Fig. 1, the outermost base pairs of bands a,b,c, and d form such a chain and close a pseudoloop. Sec. Definitions (Appendix) summarizes further definitions.

### Energy Model

To assess the energy of RNA structures, we distinguish different types of structural elements, called *loops*, i.e. hairpin loops, stacks, bulges, interior loops, or multiloops. Loops are generally defined by their outer and potentially inner closing base pairs [34].

Nearest neighbor energy models define the free energy *E*(*G*) of a secondary structure *G* as the sum of the energies of its loops *E*(*G*) = Σ _*L*∈*G*_ *E*^loop^(*L*). A prominent example is the Turner 2004 energy model [46] for pseudoknot-free RNAs, which is used by RNAfold. For pseudoknotted RNAs, Dirks and Pierce introduced the DP03 energy model, used for pseudoknot prediction in NUPACK. DP03 extends the Turner model by adding penalties for pseudoknots and bands, as well as parameters to score multiloops that ‘span’ a band [15].

#### CParty’s energy model and Vienna RNA based implementation

In CParty, we utilize the DP09 energy parameters of HotKnots 2.0, which improve upon the DP03 energy model due to training on known pseudoknotted structures [4]. Specific parameters and loop energy functions are provided in Supplementary Table 1. While HFold calculates loop energies based on Simfold [3], CParty uses the Vienna RNA library [26]. For this purpose, the energy model parameters were translated to a compatible format, allowing for better interoperability and comparability with the Vienna RNA package. Additionally, our CParty implementation supports hard constraints that restrict the partition functions to structures that leave specified bases unpaired.

**Table 1.**
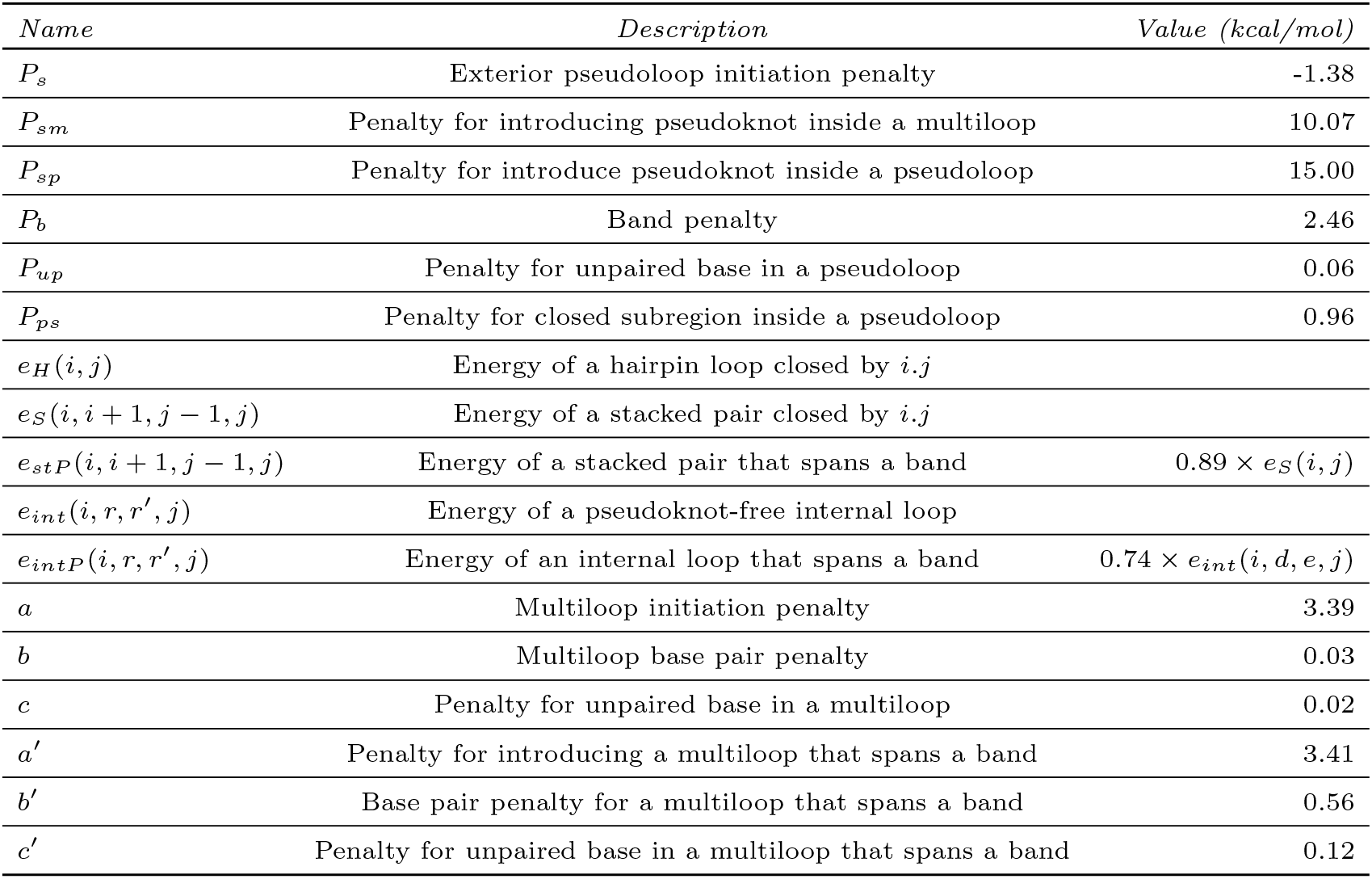
ENERGY PARAMETERS. All parameters were derived at 37 degrees celsius and 1 M salt (NaCl) concentration or extrapolated from experimental values cf. [22, 4, 35].

| Name | Description | Value (kcal/mol) |
| --- | --- | --- |
| $P_s$ | Exterior pseudoloop initiation penalty | -1.38 |
| $P_{sm}$ | Penalty for introducing pseudoknot inside a multiloop | 10.07 |
| $P_{sp}$ | Penalty for introduce pseudoknot inside a pseudoloop | 15.00 |
| $P_b$ | Band penalty | 2.46 |
| $P_{up}$ | Penalty for unpaired base in a pseudoloop | 0.06 |
| $P_{ps}$ | Penalty for closed subregion inside a pseudoloop | 0.96 |
| $e_H(i, j)$ | Energy of a hairpin loop closed by $i, j$ | |
| $e_S(i, i+1, j-1, j)$ | Energy of a stacked pair closed by $i, j$ | |
| $e_{stP}(i, i+1, j-1, j)$ | Energy of a stacked pair that spans a band | $0.89 \times e_S(i, j)$ |
| $e_{int}(i, r, r', j)$ | Energy of a pseudoknot-free internal loop | |
| $e_{intP}(i, r, r', j)$ | Energy of an internal loop that spans a band | $0.74 \times e_{int}(i, d, e, j)$ |
| $a$ | Multiloop initiation penalty | 3.39 |
| $b$ | Multiloop base pair penalty | 0.03 |
| $c$ | Penalty for unpaired base in a multiloop | 0.02 |
| $a'$ | Penalty for introducing a multiloop that spans a band | 3.41 |
| $b'$ | Base pair penalty for a multiloop that spans a band | 0.56 |
| $c'$ | Penalty for unpaired base in a multiloop that spans a band | 0.12 |

### Problem statement: partition functions over density-2 structures

Given an RNA sequence *S*, a pseudoknot-free secondary RNA structure *G*, CParty computes the hierarchically constrained partition function (CPF)

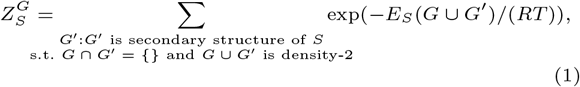

where *T* denotes the temperature (e.g, *T* = 37 ^°^C) and *R* denotes the universal gas constant (*R* ≈ 1.987 cal K^−1^ mol^−1^).

Analogous to the pseudoknot-free partition function developed by McCaskill [31], this partition function is defined as the sum of *Boltzmann weights* exp(−*E*_*S*_(*Ĝ*)*/*(*RT*)) of RNA structures *Ĝ*, where *E*_*S*_ computes the RNA energy. Extending this result, our constrained partition function sums over all density-2 structures that are the union of a given (constrained) structure *G* and a secondary structure *G*^′^. The energy *E*_*S*_ is evaluated using a pseudoknot energy model (specifically, DP09). Note that this is a true generalization, reducing to the pseudoknot-free partition function of McCaskill when the structure *G* is empty.

Boltzmann weights, *B*(*e*) := exp(−*e/*(*RT*)), and partition functions have several immediate applications in the description of the potential structures (called *ensemble*) of an RNA at equilibrium. For example, we obtain the *conditional equilibrium probability* of each structure 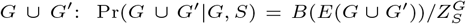, and the *ensemble free energy* 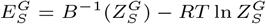 of the constrained ensemble.

### The HFold algorithm and its ambiguity

HFold efficiently minimizes the free energy over all density-2 structures *G* ∪ *G*^′^ that are hierarchically constrained by a given pseudoknot-free structure *G*. Like *G, G*^′^ must be pseudoknot-free. Energies are defined by a D&P pseudoknot energy model for the given sequence *S*. As a dynamic programming (DP) algorithm, HFold can be fully defined in terms of its recurrences.

HFold computes the total minimum free energy (MFE) as the entry *W* (1, *n*) of its DP matrix *W*, where *W* (*i, j*) denotes the MFE of the subsequence *s*_*i*_*s*_*i*+1_ … *s*_*j*_. Each *W* (*i, j*) is computed using HFold’s *W* -recurrence with the help of additional DP matrices. These matrices store MFEs under specific conditions: for example, *V* (*i, j*) is the MFE over “closed” structures that pair *i* and *j, WMB*(*i, j*) requires that *i* and *j* are the ends of a *pseudoloop*, and *VP* (*i, j*) is the MFE over the loop closed by *i*.*j* that spans a band.

Compared to pseudoknot-free prediction algorithms, HFold requires a large number of helper matrices to decompose density-2 structures and optimize correctly in the DP09 model. For instance, it distinguishes pseudoloops with the rightmost band in *G*^′^ (*WMB*^′^(*i, j*)), bands in *G* (*BE*), parts of a multiloop (*WI*), and parts of a multiloop that span a band (*WI*^′^).

#### Ambiguity

Several of HFold’s recurrences are non-trivially ambiguous, preventing a direct translation of the HFold recurrences for CParty. A good example is the decomposition of multiloops spanning a band in the *VP* recurrence, as illustrated in Figure 2.

**Fig. 2.**
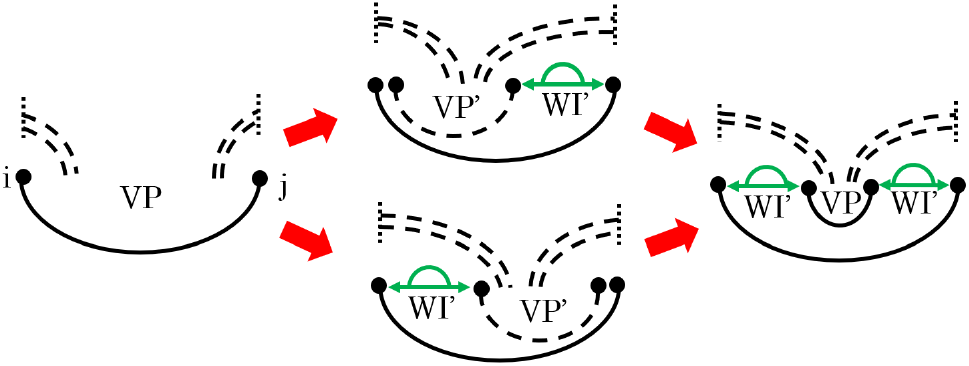
The ambiguity of computing *VP* (*i, j*) in HFold. Since *i*.*j* is a base pair of a band, it can be crossed by other bands to the left or right (dashed arcs). To handle cases where *i*.*j* closes a multiloop that spans the band, HFold utilizes two ambiguous recursion cases (middle) [22] to allow further multiloop branches on the left and/or right of the next base pair of the band. These different cases can converge (right) and produce the same structures in different ways, leading to ambiguity.

### Dataset

For the time and space complexity analysis of CParty we obtained 2808 sequences from the RNAstrand V2.0 database [2]. The smallest sequence has a length of 8 nucleotides, while the largest is 1500 nucleotides long. For each sequence, we identified the 20 most stable stem-loop by calculating only hairpin and stacking base pair energies across the whole sequence. These stem-loop structures were then used as constraints.

To assess the impact of constraint variation on CParty, we obtained 4 sequences of length 968 from the RNAstrand V2.0 database [2]. We generated 24 dinucleotide-shuffled versions for each sequence, resulting in a total of 100 sequences, using the MEME suite [6]. Using the same method as in the time and space complexity analysis, we generated the 20 most stable stem-loops for each sequence to be used as constraints, as well as the output of RNAFold, for a total of 21 input constraints for each sequence.

## The CParty Algorithm

To address the partition function problem corresponding to HFold’s energy minimization problem, we build on HFold’s decomposition of the constrained density-2 structure space. However, the ambiguity in HFold’s decomposition prevents a straightforward rewriting of the energy minimization recurrences into correct partition function recurrences by simply swapping the minimization algebra (min, +) with the ‘partition function algebra’ (+, ·). Therefore, as our core contribution, we resolve all these ambiguities by carefully rewriting HFold’s recurrences and introducing new structure classes and recurrences. This enhancement ensures a complete and unambiguous decomposition of the density-2 class of structures.

Here, we discuss the main recurrences of the CParty algorithm and refer readers to the Appendix for a detailed explanation of the remaining recurrences.

### General density-2 structures

Corresponding to the *W*(*i, j*) recurrence in HFold, *Z*_*W*_ (*i, j*) denotes the partition function over all density-2 structures 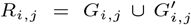 for the subsequence *s*_*i*_ … *s*_*j*_ and input substructure *G*_*i*,*j*_, taken over all choices of 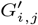.

Call a base *r covered by G*, write isCovered(*G, r*), iff it is *covered* by some base pair *k*.*𝓁* ∈ *G*, i.e. *k < r < 𝓁*. Note that *Z*_*W*_ (*i, j*) is defined only for *weakly closed* regions, where no base in the region [*i, j*] pairs with a base outside of the region. For empty region (*i > j*), *Z*_*W*_ (*i, j*) = 1—accounting for the empty structure with energy 0. Moreover, *Z*_*W*_ (*i, j*) = 0, if *i* or *j* is covered by *G*.

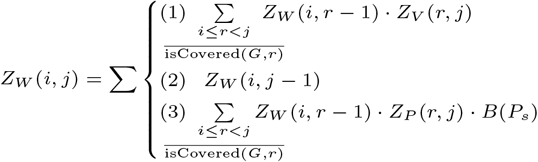

Figure 3 illustrates the three cases of *Z*_*W*_. Case (1) decomposes the structures, where *j* is paired to some *k* in [*i, j*]; it recurses to *Z*_*V*_ (*i, r*), the partition function over all structures closed by *r*.*j*. Case (2) handles structures where *j* is unpaired. Case (3) is analogous to Case (1), but *r* and *j* are left and right ends of a pseudoloop. The case recurses to *Z*_*P*_ (*r, j*) (see below), and penalizes the pseudoknot initiation (*P*_*s*_).

**Fig. 3.**
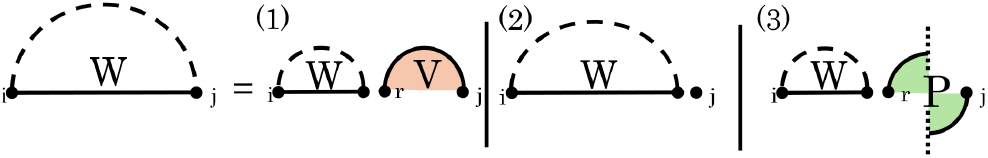
*Z*_*W*_ (*i, j*) recurrence in graphical notation: Dashed arcs indicate possible structure, each solid arc represents a base pair. The dotted vertical line indicates an overlapping chain of bands of arbitrary length and that the chain can begin or end via either *G* (above horizontal line) or *G*^′^ (below horizontal line). Filled in circles show regions covered by specific structure classes, orange for *Z*_*V*_, and green for *Z*_*P*_.

### Structures closed by a pseudoloop

The partition function over [*i, j*] where *i* and *j* are ends of a pseudoloop is calculated as *Z*_*P*_ (*i, j*). The decomposition splits of the rightmost band of the pseudoloop with ends *i* and *j*. The band can be in either *G* or *G*^′^. We handle the former case in the recurrence of *Z*_*P*_ and the latter in *Z*_*PG*_′.

Figure 4 illustrates cases of the *Z*_*P*_ recurrence. The vertical dashed line in the figure symbolizes a series of crossing alternating bands of unspecified length.

**Fig. 4.**
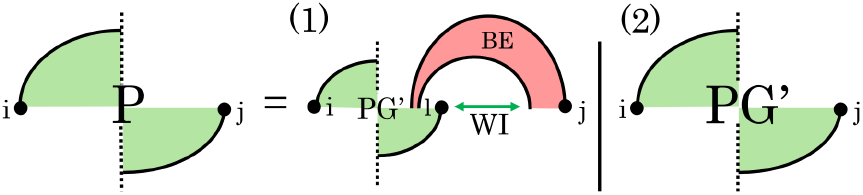
Cases of *Z*_*P*_. (1) *j* is paired in *G* and there must be some base, *l*, between *bp*_*G*_(*j*) and *j* that is paired in *G*^′^. (2) *j* is not paired in *G*, then move directly to *Z*_*PG*_′. Filled in circles show regions covered by specific structure classes, red for *Z*_*BE*_, and green for *Z*_*P*_ and *Z*_*PG*_′.

We distinguish whether *j* is paired in *G* (Case 1) or in *G*^′^ (Case 2). In Case 1, each valid structure must contain a base pair in *G*^′^ that crosses *bp*_*G*_(*j*), where *bp*_*G*_(*j*) denotes the base pair of *j* in *G*. This forms part of the pseudoloop. We consider all possible choices for the right end of this base pair, denoted as *l*. Each *l* determines unique inner and outer base pairs of the rightmost band [22].

Note that for a given *G*, only one case can be applicable (depending on whether *j* is paired in *G*). To maintain unambiguity, the corresponding sets of structures for different *l* must be disjoint, which is true for density-2 structures. Each single entry of *Z*_*P*_ is computed in linear time, and there is a quadratic number of entries.

### Pseudoknotted structures with rightmost band in *G*^′^

The partition function of the structures closed by a pseudoloop with ends *i* and *j* and rightmost band in *G*^′^ is calculated in *Z*_*PG*_′ (Fig. 5).

**Fig. 5.**
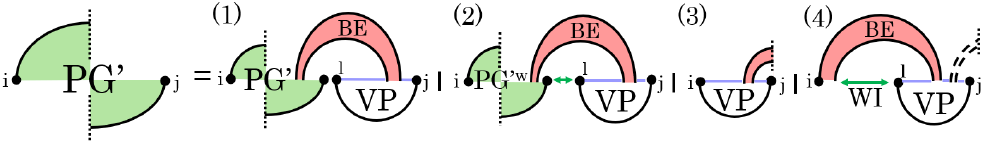
Cases of *Z*_*PG*_′. (1) handles two rightmost elements of the chain and continues. (2) is similar to (1) except there is a weakly closed region between the bands, this will be handled by *Z*_*PG*_′*w* structure class to preserve the cubic time complexity. For the end cases we have (3) leftmost band of chain in *G*^′^; and (4) leftmost band in *G*. Dashed arcs indicate possible structure, each solid arc represents a base pair. Filled in circles show regions covered by specific structure classes, green for *Z*_*PG*_′*w*. Colored lines correspond with structure classes that may or may not have any substructures: *Z*_*WI*_ in green, and purple for *Z*_*VP*_.

In case (1), *j* pairs with *l* such that *l*.*j* crosses a band of *G. Z*_*VP*_ (*l, j*) accounts for the contribution of region closed by *l*.*j*, and *Z*_*BE*_ accounts for the contribution of the band in *G*. We then recurse back to *Z*_*PG*_′ to consider the contribution of the rest of the structure. Case (2) is similar to case (1) with the only difference being the nested substructures allowed between the bands, which is handled by *Z*_*PG*_′*w* in this case. The introduction of *Z*_*PG*_′*w* (*i, j*) prevents multiple adjacent weakly closed subregions in the pseudoloop.

Cases (3-4) of *Z*_*PG*_′ are end cases, where only one or two bands, respectively, need to be accounted for. If *i* ≥ *j, Z*_*PG*_′ = *Z*_*PG*_′*w* = 0.

### Structures closed in G’, crossing G

*Z*_*VP*_ (*i, j*) is the partition function over all structures *R*_*i*,*j*_ in which *i*.*j* ∈ *G*^′^ and crosses a base pair in *G* (Fig. 6). If *i* ≥ *j, i* or *j* is paired in *G*, or *i*.*j* does not cross any base pair of *G*, then *Z*_*VP*_ (*i, j*) = 0.

**Fig. 6.**
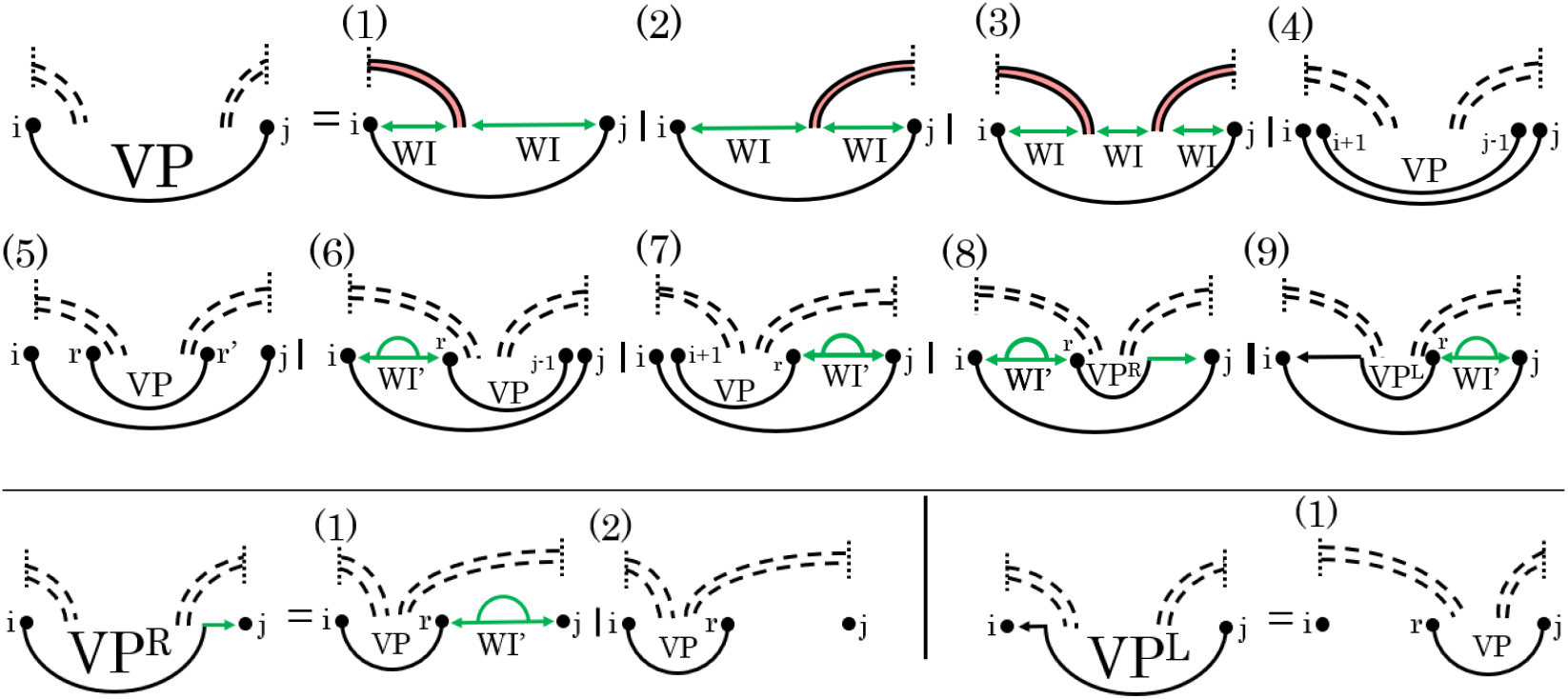
Cases of *VP*, *VP*^*R*^, and *VP*^*L*^. Top: *VP* (1 − 3) either two or three *WI* subregions (green) between *i* and *j*, band regions excluded. (4 − 5), stacked pair and internal loop, respectively. (6 − 9), *i*.*j* closes a multiloop spanning a band. Bottom-left: *VP*^*R*^, i.e., *i*.*bp*(*i*) in *G*^′^ crosses base pair in *G, bp*(*i*) ≠ *j. VP*^*R*^ (1) weakly closed non-empty region [*r* + 1, *j*], (2) empty region [*r* + 1, *j*]. Bottom-right: *VP*^*L*^, i.e., *bp*(*j*).*j* in *G*^′^ crosses base pair in *G, bp*(*j*) ≠ *i. VP*^*L*^ (1) empty region [*i, r* − 1]. Dashed arcs indicate possible structure, each solid arc represents a base pair. Colored lines correspond with structure classes: *Z*_*W I*_ in green may or may not have any substructure, but for *Z*_*W I*_′ which also has a green arc, there must be some substructure.

Cases (1-3) of *Z*_*VP*_ (*i, j*) handle nested substructures where there are no other base pairs in [*i, j*] that cross the same band(s) that *i*.*j* crosses. These nested substructures are managed by the *WI* recurrence (see Appendix). The three cases are disjoint: either *i* is covered in *G* (Case 1), *j* is covered in *G* (Case 2), or both are covered (Case 3). In Case (4), *i*.*j* and (*i* + 1).(*j* − 1) form a stacked pair; substructures created by (*i* + 1).(*j* − 1) are handled recursively by *Z*_*VP*_. In Case (5), *i*.*j* and *r*.*r*^′^ close an internal loop, and we recurse back to *Z*_*VP*_ (*r, r*^′^) for the structures formed by *r*.*r*^′^. Cases (6-9) handle *i*.*j* closing a multiloop that spans a band. In these cases, one band of the multiloop crosses the same band in *G* that *i*.*j* crosses, and the rest of the multiloop bands and unpaired bases are handled by *WI*^′^ recurrences as nested substructures. In Case (6), *r*.(*j* − 1) crosses the base pair in *G* that *i*.*j* crosses, and [*i* + 1, *r* − 1] is a non-empty weakly closed region. In Case (7), (*i* + 1).*r* crosses the base pair in *G* that *i*.*j* crosses, and [*r* + 1, *j* − 1] is a non-empty weakly closed region. In Case (8), [*i* + 1, *r* − 1] is a non-empty weakly closed region, *r*.*bp*(*r*) crosses the base pair in *G* that *i*.*j* crosses, and [*bp*(*r*) + 1, *j* − 1] is weakly closed. We introduce 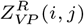 (see the bottom left part of Fig. 6), the partition function over all structures such that *i*.*r* ∈ *G*^′^ crosses a band in *G*, and *r* ≠ *j* (distinct from Case 6). Finally, in Case (9), [*r* + 1, *j* − 1] is a non-empty weakly closed region, *bp*(*r*).*r* crosses the base pair in *G* that *i*.*j* crosses, and [*i* + 1, *bp*(*r*) − 1] is empty. We introduce 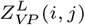 (see the bottom right part of Fig. 6), the partition function over all structures such that *r*.*j* ∈ *G*^′^ crosses a base pair in *G, r* ≠ *i* (distinct from Case 7), and [*i, r* − 1] is empty.

## Correctness

In the following, we argue that the cases of *Z*_*W*_ fully decompose the density-2 structure class, and are unambiguous. The proof sketch for correctness works by structural induction, showing the correctness of each case.

### Theorem 1

*The recurrence of Z*_*W*_ (*i, j*) *is complete, correct, and unambiguous*.

Recall that *Z*_*W*_ (*i, j*) is the partition function over the set of structures 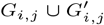 for the subsequence *s*_*i*_ … *s*_*j*_ taken over all choices of 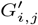 (which is pseudoknot-free, disjoint from *G*_*i*,*j*_, and such that 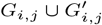 is density-2).

By definition of density-2 there are three possible cases, Case (1): *j* pairs with *r, i* ≤ *r < j*, such that *r*.*j* closes a pseudoknot-free loop, Case (2): *j* is unpaired, or Case (3): *j* is the rightmost end of a chain of crossing base pairs. These cases are disjoint; additionally, if *j* is paired and closes a pseudoknot-free loop, it cannot also be paired in the rightmost band of a pseudoloop. Therefore, the recurrence is unambiguous. Since every density-2 structure falls into one of these three cases, the *Z*_*W*_ (*i, j*) recurrence is complete. Finally, it is correct, since partition functions can be correctly inferred from smaller subproblems (which are correct by induction hypothesis). ■

Similarly we have constructed each recurrence to be complete and unambiguous by construction. Of particular importance are Cases (6 − 9) of *Z*_*VP*_ that handle a multiloop that spans a band. For a complete decomposition that preserves the *O*(*n*^3^) time complexity, 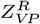 and 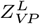 are introduced asymmetrically such that there is only one possible path to reach each structure. For example, *Z*_*VP*_ Case (8) enforces a structure somewhere in the region between *i* and *r*, and moving to 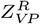 Case (1) enforces an additional structure in the subregion adjacent to *j*. To compare with *Z*_*VP*_ Case (9), similarly we enforce a structure somewhere in the region between *r* and *j*, but moving to 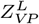 there is no possible case to introduce an additional structure adjacent to *i*. Thus, we avoid any ambiguity in *Z*_*VP*_ decomposition.

## Complexity

Starting with the *Z*_*W*_ recurrence, we observe that its time and space complexity depend on those of *Z*_*V*_ and *Z*_*P*_. Since *Z*_*V*_ handles pseudoknot-free loops, its time complexity is *O*(*n*^3^), and its space complexity is *O*(*n*^2^), where *n* is the length of the input sequence.

*Z*_*P*_ deals with pseudoloops. As *Z*_*P*_ matches the *WMB* recurrence of HFold, and HFold has been proven to have time and space complexities of *O*(*n*^3^) and *O*(*n*^2^) respectively, the same applies to *Z*_*P*_. We further empirically verify *Z*_*P*_ ‘s time and space complexity (see Empirical Results). Therefore, *Z*_*W*_ ‘s time and space complexity remain *O*(*n*^3^) and *O*(*n*^2^), respectively.

Similarly, all other cases remain within the *O*(*n*^3^) time and *O*(*n*^2^) space complexity. For example, the time complexity for both *Z*_*PG*_′ and *Z*_*PG*_′*w* is *O*(*n*^3^), as both cases involve searching over all values of *l* for a given region [*i, j*]. The time complexity of *Z*_*VP*_ is dominated by the search over the region [*i, j*] to find the value of *r*, which is also *O*(*n*^3^).

## Empirical Results

Since CParty solves the conditional partition function for density-2 structures for the first time, it cannot be directly *benchmarked* against existing algorithms. Nevertheless, some comparisons to RNAfold and NUPACK remain meaningful and can provide insights.

Recall that in the special case of an empty input structure *G*, CParty computes a *pseudoknot-free partition function Z*_pkfree_. As plausibility check, we first compared the ensemble free energy computed by CParty for *Z*_pkfree_ to the ensemble free energy for pseudoknot-free structures computed by RNAFold [26]. Here, CParty perfectly reproduces the results of RNAfold (Fig. 7a), using Turner2004 parameters [29] without dangle energies.

**Fig. 7.**
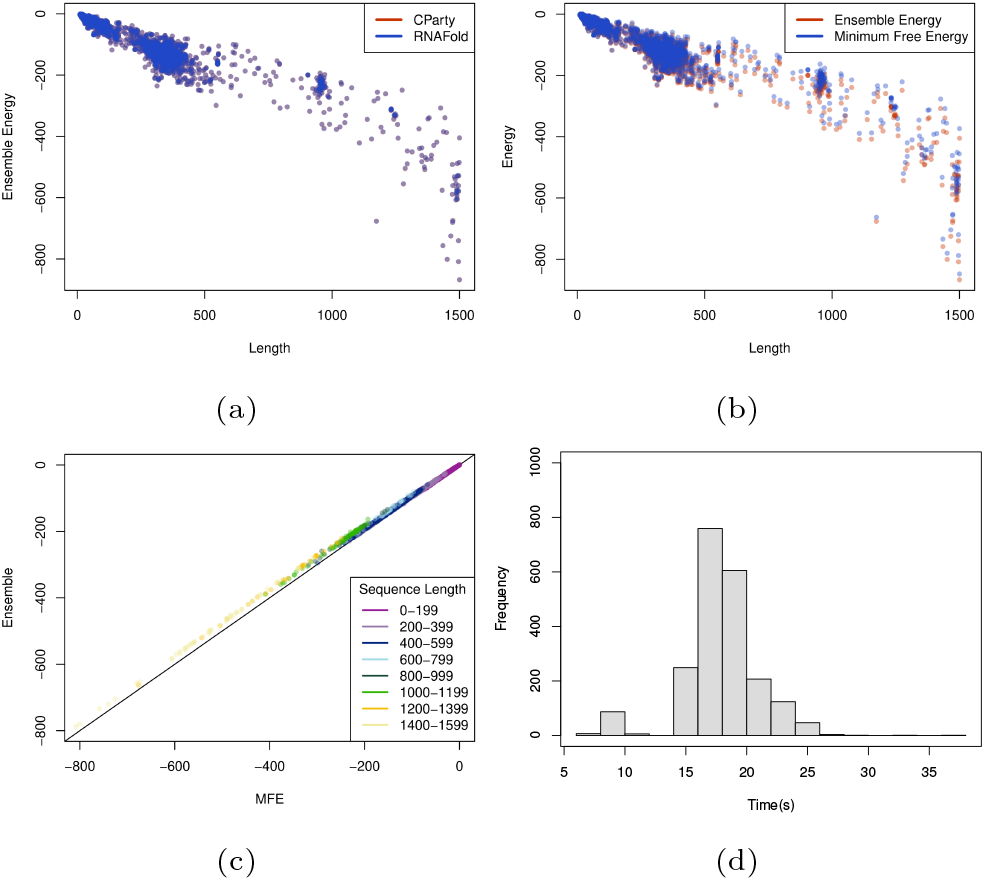
We considered 2733 sequences with length up to 1500 analyzed from the RNAstrand V2.0 database [2]. (a) Ensemble free energies without constraints via CParty (shown in red) and RNAfold (shown in blue). General agreement is observed between the two. (b) Ensemble free energies vs minimum free energy of CParty with constraints (pseudoknotted). (c) Each sequence is plotted as its ensemble energy from CParty vs the minimum free energy from HFold. Colors represent the lengths of the sequences. A diagonal line represents a 1 to 1 for ensemble energy to minimum free energy. (d) We plot the results of CParty given a sequence and an input structure. We took 4 sequences of equal length from the RNAStrand database [2] and created 24 dinucleotide shuffled versions for each of them. Each sequence had 21 varying input structures; time was placed in a histogram to show the distribution given different inputs.

We then sought to assess the empirical time and space of computing the CParty partition function, *Z*, against RNAFold and NUPACK. We chose RNAfold as a benchmark for our lower bound and NUPACK as it is the only pseudoknotted partition function calculation algorithm. Since CParty requires an input structure in addition to the RNA sequence, for each sequence we identified the stem-loop structure with the lowest free energy, as detailed in Section Dataset, and used it as input to CParty and RNAFold (NUPACK’s algorithm does not accept a partial structure as input), for a fair runtimes comparison. All experiments were performed on the Digital Research Alliance of Canada’s Cedar cluster. We measured runtime using user time (see Fig. 8a), and memory using maximum resident set size (see Fig. 8b). The maximum time and memory used by CParty was 103.42 seconds and 117212 KB. In comparison, RNAFold had a maximum time of 36.39 seconds and 52208 KB. The expected increase in time and space usage when transitioning from pseudoknot-free to pseudoknotted structures in CParty is due to the need for new data structures and additional recurrence relations. As NUPACK requires a large amount of memory, its results were limited to sequences of max length 100. The maximum time and space for NUPACK on this subset of our dataset were 23.05 seconds and 460908 KB (see Fig. 8a and 8b in blue). As seen in Figure 8, CParty’s time and space complexities closely match those of RNAFold.

**Fig. 8.**
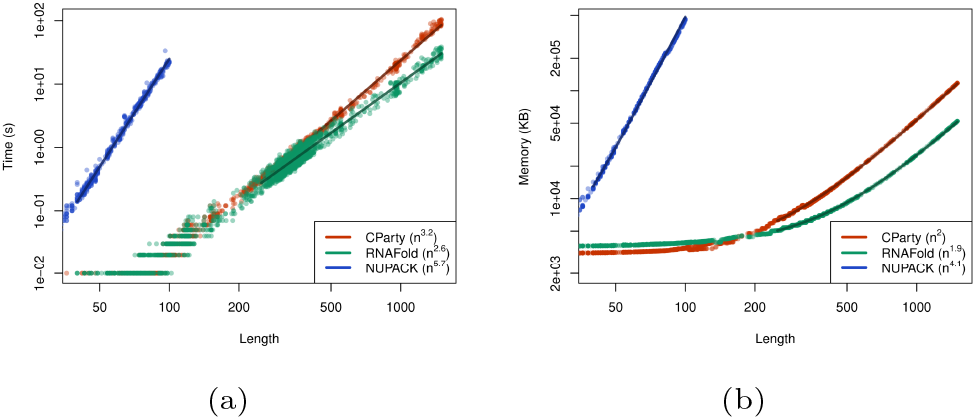
Time and space consumption of CParty vs. RNAFold and NUPACK on our dataset, when given an RNA sequence and a pseudoknot-free structure as input. (a) Memory Usage (maximum resident set size in KB) versus length (log-log plot) over all benchmark instances. The solid line shows an asymptotic fit (*c*_1_ + *c*_2_*n*^*x*^) for sequence length *n*, constants *c*_1_,*c*_2_, and exponent x for the fit. We ignored all values < 250 for CParty and RNAFold and all values < 40 for NUPACK. (b) Run-time (s) versus length (log-log plot) over all benchmark instances. For each tool in both plots, we report (in parenthesis) the exponent *x* that we estimated from the benchmark results; it describes the observed complexity as Θ(*n*^*x*^).

To assess the potential impact of the input structure on the performance of CParty, we calculated the constrained partition function on 100 sequences of length 968 with a total of 2100 various input structures, as detailed in Section Dataset. As shown in Figure 7d, little variation is observed in memory given different input structures with the 25th and 75th percentiles showing a difference of 10 KB. Figure 7d also provides a median time of 18 seconds with the 25th and 75th percentiles showing a difference of only 2 seconds. While variations in CParty’s time and space usage are expected, those were not deemed significant.

### Analysis of SARS-CoV-2 frameshift structure

There has been extensive research into predicting structure of the SARS-CoV-2 frameshift sequence, which includes both computational efforts [41, 45, 44] and experimental probing experiments [19, 28, 50, 51]. The frameshift sequence is believed to form a density-2 pseudoknotted structure [24, 40].

Employing CParty with different fixed input structures, here we provide a stochastic view of suboptimal structures for the SARS-CoV-2 frameshift stimulating structure ensemble. Combining the available SHAPE reactivity probing datasets and various thermodynamic-based algorithms, we previously identified the top-most energetically favourable initial stems for the SARS-CoV-2 77 nucleotide frameshift pseudoknot sequence [41, 45, 44]. Here, we utilize the top two stems (referred to as initial stem 1 and 2) to explore the structural ensemble for the frameshift sequence. These two stems were identified as pivotal for formation of two of the main structural motifs, referred to as 3 − 3 and 3 − 6 [40](see Fig. 10b).

Following the pipeline of Fig. 9, with each of the two initial stems as constraint, we employ CParty to compute ensemble free energy for sequences of decreasing length (taking 7 bases one at a time from the 5’ end), to simulate the effects of the translocating ribosome [5, 14].

**Fig. 9.**
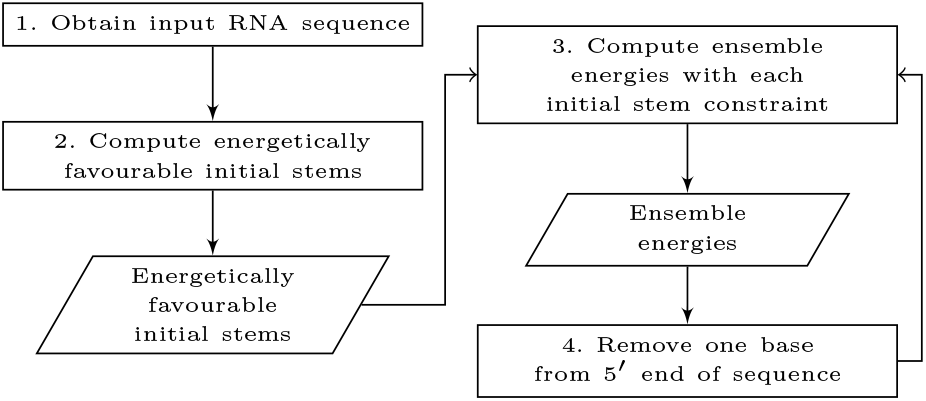
CParty constrained ensemble energy pipeline. Rectangles dictate actions, parallelograms denote outputs.

As seen in Fig. 10, ensemble free energies constrained by stems 1 and 2 are close to one another for the 77 length frameshift sequence. However, at the 5th base removal from the 5’ end (see the red rectangle in Fig. 10), the ensemble free energy for stem 2 increases, suggesting a significant change of structural ensemble at this point. We further investigated this possible structural change using Iterative HFold [20] with frameshift sequence and initial stems 1 and 2 as input and decreasing the length by 7 bases (one base at a time) from the 5’ end. We noticed that both stem 1 and 2 can form the 3 3 structural motif at original sequence length (77). However, at the marked transition form (red rectangle), the 3 6 motif (also referred to as the native structure) becomes more stable; this structural motif is only compatible with stem 1. This transition observed through both ensemble free energy change as well as structurally supports the hypothesis that destabilization of initial stem 2 *facilitates* subsequent refolding of the native-type pseudoknot [44].

**Fig. 10.**
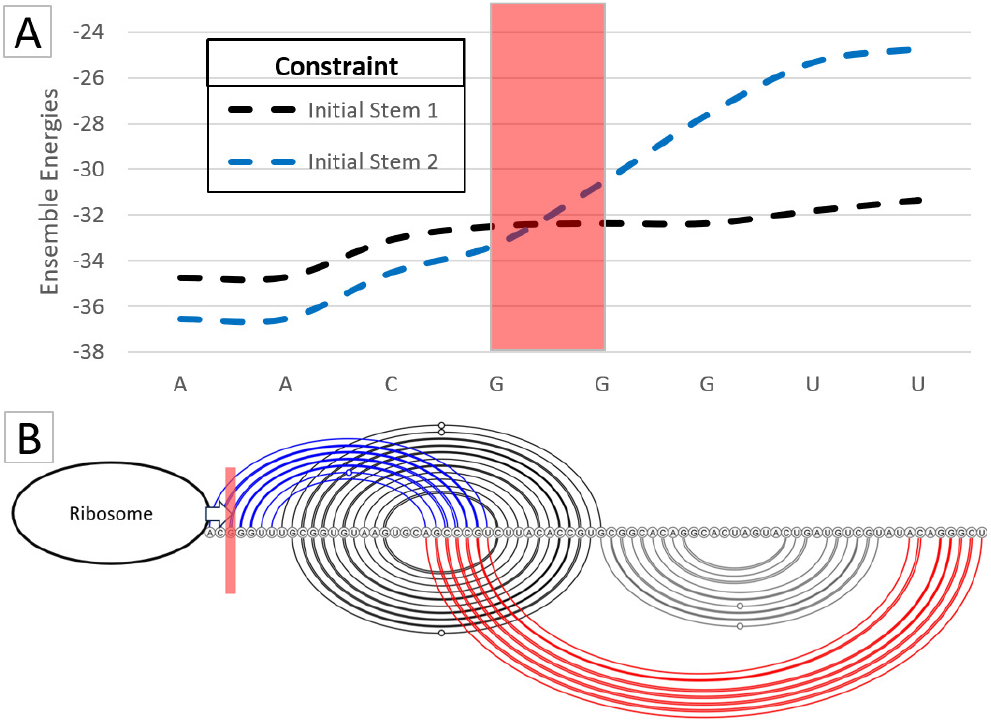
SARS-CoV-2 secondary structure motif transition. (a): Constrained ensemble energies for decreasing SARS-CoV-2 sequence lengths (decreasing from 77 to 70 nt, left to right, sequence labeled on x-axis). (b) Arcs represent base pairs. Top arc diagram: 3_3 motif [40], initial stem 1 in black, initial stem 2 in blue. Bottom arc diagram: 3_6 motif, note that this motif is *not compatible* with initial stem 2. (a & b): Red rectangles highlight the location of a transition from the 3_3 motif to the 3_6 motif. When the ribosome destabilizes the 3_3 motif base pairs (blue arcs) to the left of the red rectangle, refolding of the native-type pseudoknot (red arcs) is expected.

## Discussion

In this work, we introduce CParty, a novel biologically motivated algorithm that follows the hierarchical folding hypothesis to efficiently compute the constrained partition function (CPF) for density-2 RNA secondary structures. CParty takes an RNA sequence and a pseudoknot-free structure *G* as input and computes the CPF over all density-2 structures *G* ∪ *G*^′^, where *G*^′^ is pseudoknot-free and disjoint from *G*.

CParty was developed by addressing the ambiguities in the HFold algorithm [22]. While HFold relies on Simfold [3] for pseudoknot-free energy calculations, CParty utilizes the efficient and well-maintained ViennaRNA library [26] and supports various energy models.

CParty handles the class of density-2 structures, which includes a wide range of pseudoknots such as kissing hairpins and interleaved bands of infinite length with arbitrarily nested substructures of the same class. By employing a hierarchical folding approach, CParty achieves a runtime complexity of *O*(*n*^3^) and a space complexity of *O*(*n*^2^). We evaluated the empirical time and space usage of CParty against RNAfold and NUPACK on a large dataset of RNA sequences of varying lengths, demonstrating CParty’s efficiency in handling large RNA sequences.

Correctly identifying the input structure *G* is an important factor when using our algorithm. As noted in previous studies [20, 45], utilizing the most stable pseudoknot-free stem-loops is effective in identifying both the minimum free energy (MFE) structure and low-energy suboptimal structures—those energetically close to the MFE structure. Repeatedly sampling the hierarchical distribution with multiple fixed structure choices for *G* can help identify possible folding paths to different secondary structure motifs. Although the input structure influences our algorithm’s runtime and memory usage, we found this impact to be minimal.

In this work, we demonstrated CParty’s application in characterizing structural motifs in the SARS-CoV-2 frameshift element. We believe our algorithm can be similarly used in other structure-function characterizations and aid in the development of novel therapeutics.

Under hierarchical folding assumptions, CParty enables us to calculate the probability of observing a density-2 structure *G* ∪ *G*^′^ at equilibrium for an RNA *S* as the product of the pseudoknot-free probability of *G* (following McCaskill’s method) and the conditional probability Pr(*G* ∪ *G*^′^|*G, S*).

Building on this concept, CParty supports sampling from the corresponding hierarchical structure probability distribution. While we plan to study hierarchical sampling explicitly in future work, it can be achieved through direct stochastic traceback from CParty’s dynamic programming matrices. This process involves a two-step approach: first, sampling pseudoknot-free structures *G* [13], and then drawing from the hierarchical distribution constrained by *G*.

By leveraging these capabilities, CParty offers a powerful and efficient method for exploring RNA secondary structures, paving the way for further advancements in RNA research.

## Author contributions statement

L.T., S.W., Y.P., U.S. and H.J. conceptualized the work. M.G. implemented the algorithm, and M.G. and L.T. conducted the experiments and analysis. All authors wrote and reviewed the manuscript.

## Acknowledgments

We thank and acknowledge the Computational Biology Research and Analytics Lab for invaluable feedback.

## Appendix

### Definitions

- **RNA molecule**: A sequence of nucleotides, or bases, of length *n*, of which there are four types: Adenine (A), Guanine (G), Cytosine (C), and Uracil (U).
- **Base pair**: When a RNA folds, bonds form between the bases of the molecule, where each base may pair with at most one other base.
- **RNA structure R**: The set of base pairs *i*.*j*, 1 ≤ *i < j* ≤ *n* such that no index occurs in more than one base pair, and each base pair is one of the canonical base pairs: {*A* − *U, C* − *G, G* − *U*}.
- **bp**_**R**_(**i**): We let *bp*_*R*_(*i*) denote the index of the base that is paired with base *i* in *R*, if any.
- **cross**: if *i*.*j, i*^′^.*j*^′^, and *i < i*^′^ *< j < j*^′^, we say that the pair *i*.*j* crosses the pair *i*^′^.*j*^′^ (and *i*^′^.*j*^′^ crosses *i*.*j*)
- **Pseudoknotted base pair**: We say that i.j. is a pseudoknotted base pair if for some other base pair i’.j’ in R, i.j crosses i’.j’.
- **Pseudoknot-free structure**: If there are no pseudoknotted base pairs in the given structure, it is called pseudoknot free secondary structure.
- **Cover**: Let *G* be a pseudoknot-free structure. Base pair *i*.*j covers* base k if *i < k < j* and there is no other base pair *i*^′^.*j*^′^ where *i < i*^′^ *< k < j*^′^ *< j*.
- **is Covered(G**,**k)**: true iff some base pair of G covers k.
- **Region [***i, j*]: Sequence of indices between i and j inclusive.
- **Disjoint region**: two regions [*i, j*] and [*i*^′^, *j*^′^] are disjoint if no index is in both regions, i.e *j < i*^′^ or *j*^′^ *< i*
- **Weakly closed region**: A region is weakly closed if no base connects a base in the region to a base outside the region.
- **Closed region**: A weakly closed region with at least two bases,[*i, j*], is closed, if it cannot be partitioned into two smaller weakly closed regions. Note that if [*i, j*] is closed, then both *i* and *j* must be paired, although not necessarily with each other [34].
- **Pseudoknotted closed region**: a closed region [*i, j*] of a structure *R* such that *i*.*bp*_*R*_(*i*) and *bp*_*R*_(*j*).*j* are pseudoknotted base pairs.
- **directly banded in**: For a psuedoknotted base pair *i*.*j*, we say *i*.*j* is *directly banded in i*^′^.*j*^′^, denoted *i*.*j* ⪯ *i*^′^.*j*^′^, if *i*^′^ ≤ *i < j* ≤ *j*^′^ and [*i*^′^ + 1, *i* − 1] and [*j* + 1, *j*^′^ − 1] are weakly closed regions
- **Band**: Consider a maximal chain of ⪯. The minimum (maximum) base pair in the maximal chain is the band’s inner (outer) closing pair. If *i*.*j* is the outer and *i*^′^.*j*^′^ the inner closing pair of a band, then [*i, i*^′^] and [*j*^′^, *j*] are the band’s regions
- **Pseudoloop**: Let [*i, j*] be a pseudoknotted closed region. Then the unpaired bases and base pairs associated with [*i, j*], together with the closing base pairs of the band associated with [*i, j*], are members of a *pseudoloop*. The base pairs *i*.*bp*_*R*_(*i*) and *bp*_*R*_(*j*).*j* are the closing base pairs of the pseudoloop.
- **Bi-secondary structure**: A structure *R* such that *R* can be formed by the union of two disjoint pseudoknot-free secondary structures [49].
- **Density**: We define density as follows: Let L be a psuedoloop and *i*.*bp*_*R*_(*i*) and *bp*_*R*_(*j*).*j* be the closing base pairs of L. Let #B(L,k) be the number of bands associated with L that cross k. Then the density of L is the max #B(L,k) for all k in region [*i, j*]. The density of a structure, R, is the maximum density of L over all pseudoloops L of R. We say R is a density-2 structure if the density of R is at most 2.

We provide Fig. 11 and Fig. 12 for density-2 structure class intuition.

**Fig. 11.**
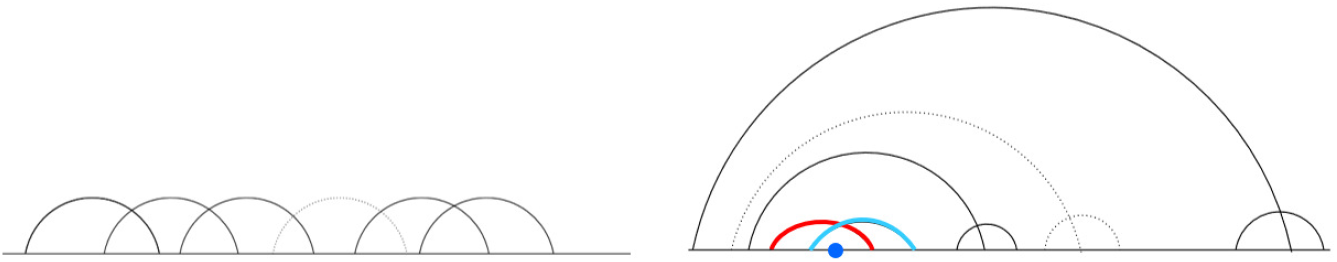
Arc diagram representation of two density-2 structures, each structure contains an arbitrary number and depth of bands [22]. Blue dot covered by red and light blue bands indicates how association with closed region maintains the density-2 property. Figure modified from [22].

**Fig. 12.**
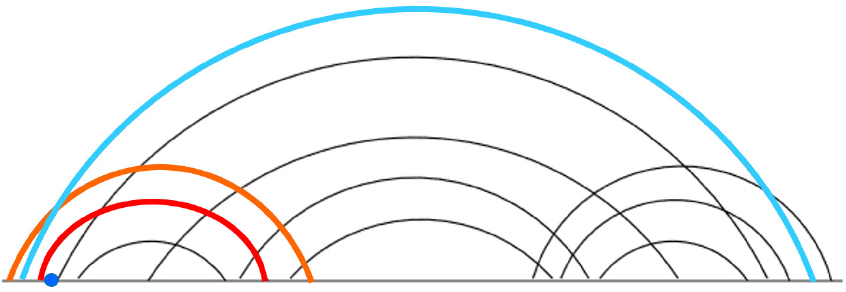
Example of a *bi-secondary structure* that is not a density-2 structure [22]. Blue dot covered by red, orange, and light blue bands indicate density-2 property not satisfied. Figure modified from [22].

### Pseudoknotted density-2 structures

Consider the class of secondary structures 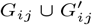 that contain pseudoknotted base pairs and cannot be partitioned into two independent substructures for two regions [*i, r*] and [*r* + 1, *j*], for some *r*. Here, we define *Z*_*P*_ (*i, j*), the partition function over [*i, j*] when [*i, j*] is a pseudoknotted closed region, containing a chain of two or more successively overlapping bands that must alternate between *G*_*i*,*j*_ and 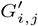, possibly with nested substructures interspersed throughout.

Note that the rightmost band of the pseudoloop may be in *G* or *G*^′^. In order to calculate the energies of substructures in such a structure in our recurrences, we use additional terms including the following: *Z*_*BE*_, *Z*_*VP*_, and *Z*_*WI*_. Roughly, these account for energies of bands spanned by base pairs of *G*_*i*,*j*_, regions enclosed by pseudoknotted base pairs of 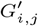 (excluding part of those regions that are within a band of *G*_*i*,*j*_), and nested, weakly closed regions, respectively.

For *Z*_*P*_ (*i, j*) base case, if *i* ≥ *j*, then *Z*_*P*_ (*i, j*) = 0, since the substructure is empty, and thus cannot be pseudoknotted. Otherwise, there are two cases (cf. Fig. 4), Case (1): *j* is paired in *G*, or Case (2): *j* is not paired in *G*. If *j* is paired in *G*, then in the MFE structure, some base *l* with *bp*_*G*_(*j*) *< l < j* must be paired in *G*^′^, causing *bp*_*G*_(*j*).*j* to be pseudoknotted. We consider all possible choices of *l*. Once *l* is fixed, the inner base pair of the band whose outer base pair is *bp*_*G*_(*j*).*j* is also determined (e.g., *b*_(*i*,*l*)_ or 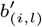 cf. [22]). The *B*(*P*_*b*_) and *Z*_*BE*_ terms in Case (1) account for band energy, a *Z*_*W I*_ term accounts for the energy of a weakly closed region that is nested in the band, and the remaining energy is represented by the *Z*_*PG*_′ term. If *j* forms a pseudoloop with *i* and is not paired in *G*, we move to *Z*_*PG*_′ Case (2), partition function over pseudoknotted density-2 structures with rightmost band in *G*^′^. Here we note these two cases are disjoint and *Z*_*P*_ is unambiguous, with *j* either paired in *G* or *G*^′^. Conditions for terms in equation summation are shown in blue.

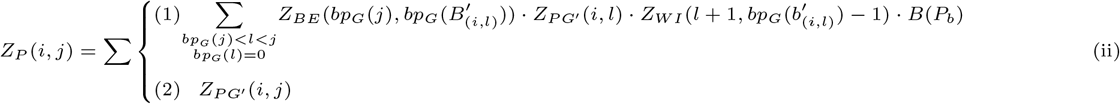

### Pseudoknotted structures with rightmost band in *G*^′^

When *i* and *j* form a pseudoloop where the rightmost band of the pseudoloop is not in *G*, i.e., *bp*_*G*_(*j*) = 0, it must be part of the *G*^′^ structure. *Z*_*PG*_′ handles energy of all structures where the rightmost band is not in *G*, but is part of the structure *G*^′^. Therefore, Cases (1 − 2) of *Z*_*PG*_′ (cf. Fig. 5) are used to account for the energy of the region spanned by the rightmost two bands using *B*(2*P*_*b*_),

*Z*_*BE*_ and *Z*_*VP*_ (*l, j*); and recursively calling either *Z*_*PG*_′, or to *Z*_*PG*_′*w* if there is a weakly closed region between the bands. To avoid allowing multiple adjacent weakly closed subregions in the pseudoloop, we must introduce *Z*_*PG*_′*w* (*i, j*), the partition function over all pseudoknotted density-2 structures between *i* and *j* where the rightmost band of the pseudoloop is part of the structure of *G*^′^ and the structure is weakly closed. For band border details see [22]. *Z*_*PG*_′ Cases (3 − 4) are end cases, where only one or two bands, respectively, need to be accounted for so no recursive call is made. For the base case when *i* ≥ *j, Z*_*PG*_′ = *Z*_*PG*_′*w* = 0.

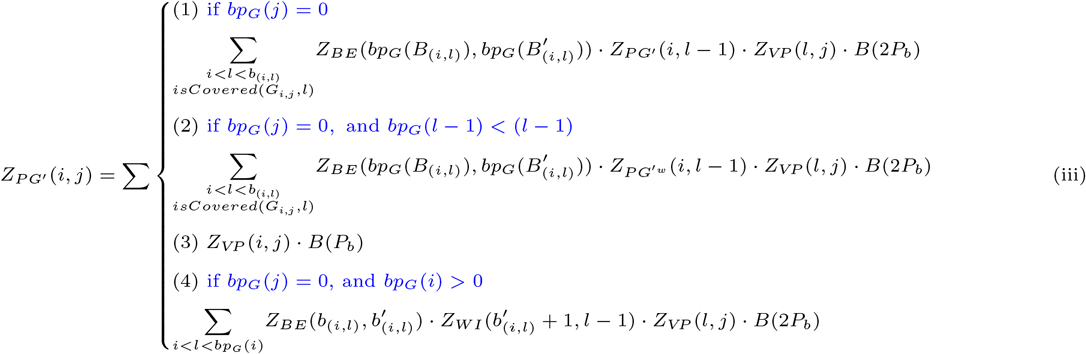

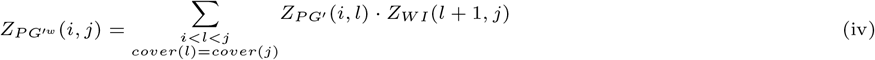

### Structures closed in G’, crossing G

*Z*_*VP*_ (*i, j*) is the partition function over all structures *R*_*i*,*j*_ in which *i*.*j* ∈ *G*^′^ and crosses a base pair in *G*. The energy of *R*_*i*,*j*_ is the energy of all loops within *R*_*i*,*j*_ that are not inside a band whose base pairs are in G and which crosses *i*.*j*. If *i* ≥ *j, i* or *j* is paired in *G*^′^, or *i*.*j* does not cross any base pair of *G*, then *Z*_*VP*_ (*i, j*) = 0, otherwise *Z*_*VP*_ (*i, j*) is computed as follows.

*Z*_*VP*_ (*i, j*) Cases (1 − 3) handle structures where there are no other base pairs in *R*_*i*,*j*_ that cross the band(s) *i*.*j* crosses. These cases are unambiguous: either *i* is covered, *j* is covered, or both. In Case (4), (*i* + 1).(*j* − 1) forms a stacked pair (with its energy computed through *e*_*stP*_). In Case (5), *i*.*j* and *r*.*r*^′^ close an internal loop (with its energy computed through *e*_*intP*_). In Case (6), *r*.(*j* − 1) crosses a base pair in *G* and [*i* + 1, *r* − 1] is a weakly closed non-empty region (multiloop spanning band initiation, base pair, and unpaired base penalties, *a*^′^, *b*^′^, and *c*^′^, respectively). In Case (7), (*i* + 1).*r* crosses a base pair in *G* and [*r* + 1, *j* − 1] is a weakly closed non-empty region. In Case (8), [*i* + 1, *r* − 1] is a weakly closed non-empty region, *r*.*bp*(*r*) crosses a base pair in *G*, and [*bp*(*r*) + 1, *j* − 1] is either empty or non-empty and weakly closed. Therefore, we introduce 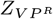 (cf. Fig. 6), the partition function over all structures such that *i*.*bp*(*i*) ∈ *G*^′^ crosses base pair in *G*, and *bp*(*i*) ≠ *j* (distinct from Case (6)). Finally, in Case (9), [*r* + 1, *j* − 1] is a weakly closed non-empty region, *r*.*bp*(*r*) crosses a base pair in *G*, and [*i* + 1, *bp*(*r*) − 1] is empty. Therefore, we introduce 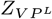, the partition function over all structures such that *bp*(*j*).*j* ∈ *G*^′^ crosses base pair in *G, bp*(*j*) ≠ *i* (distinct from Case (7)), and [*i* + 1, *bp*(*j*) − 1] is empty.

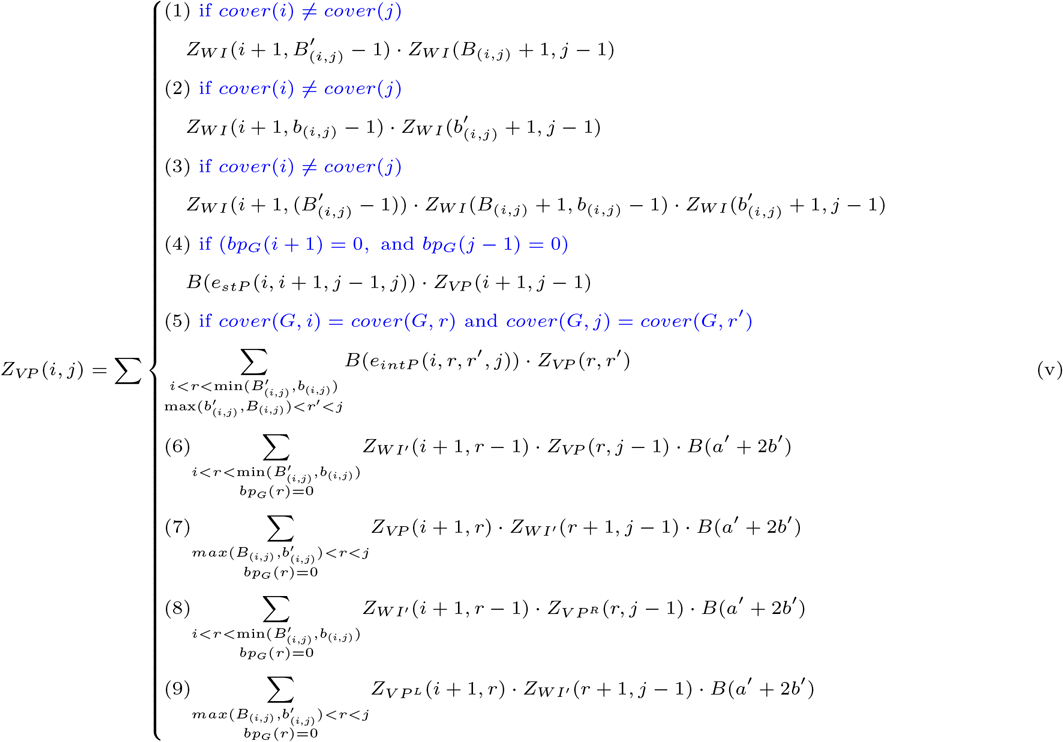

*i*.*bp*(*i*) *in G*^*′*^ *crosses base pair in G, bp*(*i*) *not equal to j*

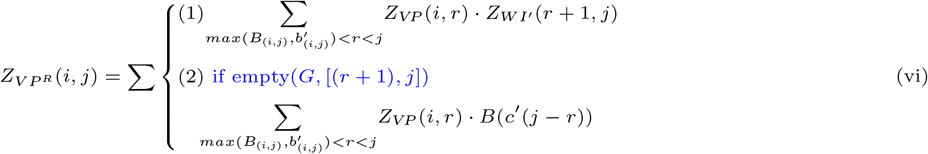

*bp*(*j*).*j in G*^*′*^ *crosses base pair in G, bp*(*j*) *not equal to i*

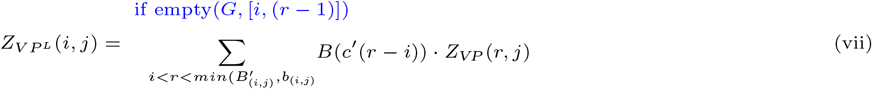

*i*.*j* closes a multiloop

*Z*_*V M*_ (*i, j*) is the partition function over all structures *R*_*i*,*j*_ for region [*i, j*], if [*i, j*] is weakly closed and *i*.*j* closes a multiloop. Otherwise, *Z*_*V M*_ (*i, j*) = 0.

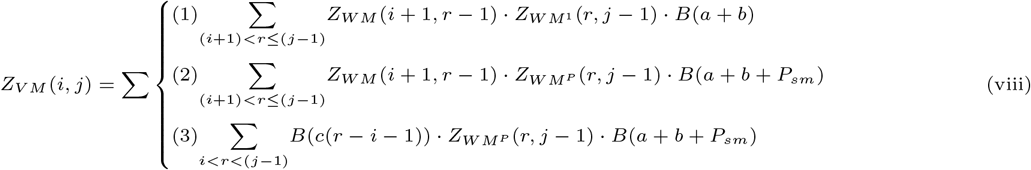

*i* and *j* on a multiloop.

*Z*_*W M*_ (*i, j*) is the partition function over all structures *R*_*i*,*j*_ for region [*i, j*], if [*i, j*] is weakly closed, not empty, and *i* and *j* are on a multiloop. For base case where *i* ≥ *j, Z*_*W M*_ (*i, j*) = 0.

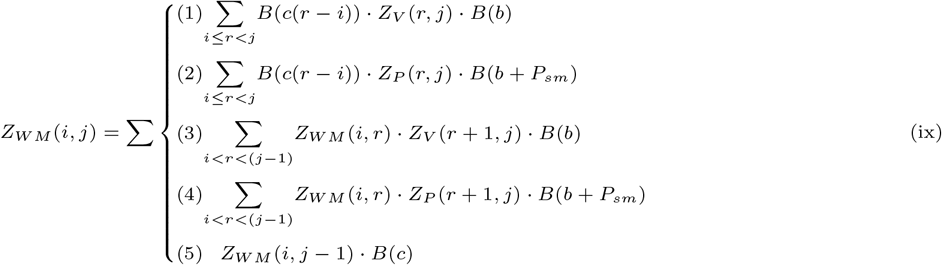

*i and j on a multiloop, terminal stem pseudoknot-free*

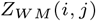 is the partition function over all structures *R*_*i*,*j*_ for region [*i, j*], if [*i, j*] is weakly closed, not empty, and *i* and *j* are on a multiloop (terminal stem, cf. Fig. 13). With Case (1), *i*.*j* form a pseudoknot-free loop, and Case (2), *j* is unpaired. For base case where 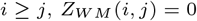.

**Fig. 13.**
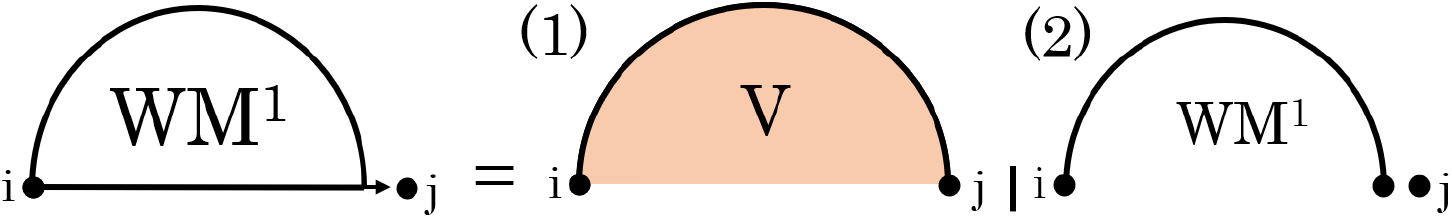
Cases of *WM* ^1^. (1) terminal stem pseudoknot-free, (2) *j* is unpaired.

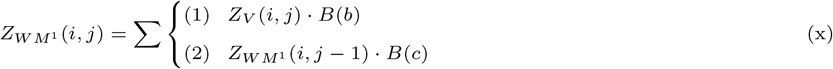

*i and j on a multiloop, terminal stem pseudoknotted*

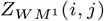 is the partition function over all structures *R*_*i*,*j*_ for region [*i, j*], if [*i, j*] is weakly closed, not empty, and *i* and *j* are on a multiloop (pseudoknotted terminal branch, cf. Fig. 14). With Case (1), *i*.*j* form a pseudoloop, and Case (2), *j* is unpaired. For base case where 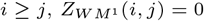.

**Fig. 14.**
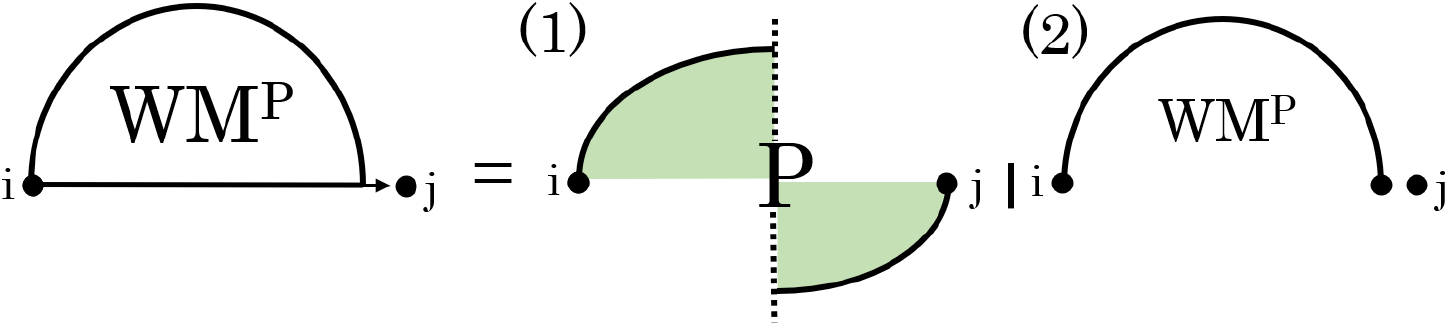
Cases of *WM*^*P*^. (1) terminal stem pseudoknotted, (2) *j* is unpaired.

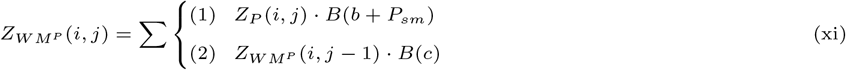

#### Weakly closed subregions inside pseudoloop

*Z*_*W I*_ (*i, j*) is the partition function over all structures *R*_*i*,*j*_ given that [*i, j*] is weakly closed (*cover*(*i*) = *cover*(*j*) ≠ 0), and *R*_*i*,*j*_ is inside a pseudoloop. If *i* = *j* and *bp*_*G*_(*i*) = 0, [*i, j*] is empty and *Z*_*W I*_ (*i, j*) = *P*_*up*_, i.e., penalty for unpaired base in a pseudoloop. *Z*_*W I*_ (*i, j*) = 0, if *i > j*. Otherwise, *Z*_*W I*_ (*i, j*) = 0 (*cover*(*i*) ≠ *cover*(*j*), subregion not weakly closed). *Z*_*WI*_ is similar to *Z*_*W*_ with additional penalties for base pair, unpaired bases, and pseudoknot initiation inside a pseudoloop.

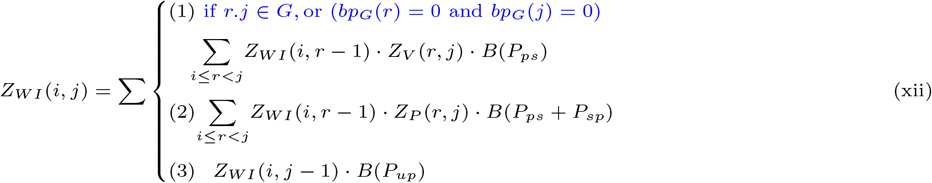

#### Non-empty weakly closed subregion inside band

*Z*_*W I*_′ (*i, j*) is the partition function over all nonempty structures *R*_*i*,*j*_, if [*i, j*] is weakly closed with respect to *G*, given that *R*_*i*,*j*_ is inside a band. Otherwise, *Z*_*W I*_′ (*i, j*) = 0. *Z*_*W I*_′ is similar to *Z*_*W I*_ with additional penalties for base pair or unpaired base in multiloop spanning a band, and pseudoknot initiation inside a multiloop.

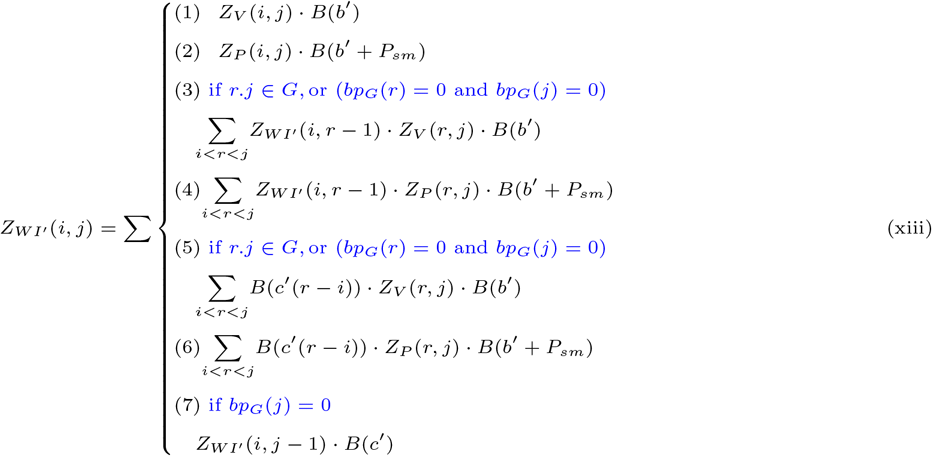

*i*.*j* close a loop

*Z*_*V*_ (*i, j*) is the partition function over all structures *R*_*i*,*j*_ for region [*i, j*], if [*i, j*] is weakly closed or empty and *i*.*j*. Otherwise *Z*_*V*_ (*i, j*) = 0. This subfunction and *Z*_*V BI*_ (*i, j*) to follow are unchanged from Mathews et al. [30] pseudoknot-free algorithm, i.e., penalties for hairpin loop, stacked base pairs, internal/bulge loop, or multiloop.

*i*.*j close an internal/bulge loop*

*Z*_*V BI*_ (*i, j*) is the partition function over all structures *R*_*i*,*j*_ for region [*i, j*], if [*i, j*] is weakly closed or empty and *i*.*j* closes a bulge or internal loop. Otherwise *Z*_*V BI*_ (*i, j*) = 0 [30].

[*i, i*^′^] ∪ [*bp*(*i*^′^), *bp*(*i*)] band region

*Z*_*BE*_ (*i, i*^′^) is the partition function over the band [*i*^′^, *i*] ∪ [*bp*_*G*_(*i*), *bp*_*G*_(*i*^′^)], if *i* ≤ *i*^′^ *< bp*_*G*_(*i*^′^) ≤ *bp*_*G*_(*i*); otherwise *Z*_*BE*_ (*i, i*^′^) = 0. In Case (1), *bp*(*i* + 1).*bp*(*i*) − 1 form a stacked pair in *G*. In Case (2), *i*.*bp*(*i*) and *l*.*bp*(*l*) in *G* close an internal loop. In Case (3), [*i* + 1, *l* − 1] and [*bp*(*l*) + 1, *bp*(*i*) − 1] are both weakly closed non-empty region. In Case (4), [*i* + 1, *l* − 1] is a weakly closed non-empty region and [*bp*(*l*) + 1, *bp*(*i*) − 1] is empty. In Case (5), [*bp*(*l*) + 1, *bp*(*i*) − 1] is a weakly closed non-empty region and [*i* + 1, *l* − 1] is empty (cf. Fig. 15). For base case, *Z*_*BE*_ (*i, i*) = 0 if *i < bp*_*G*_(*i*).

**Fig. 15.**
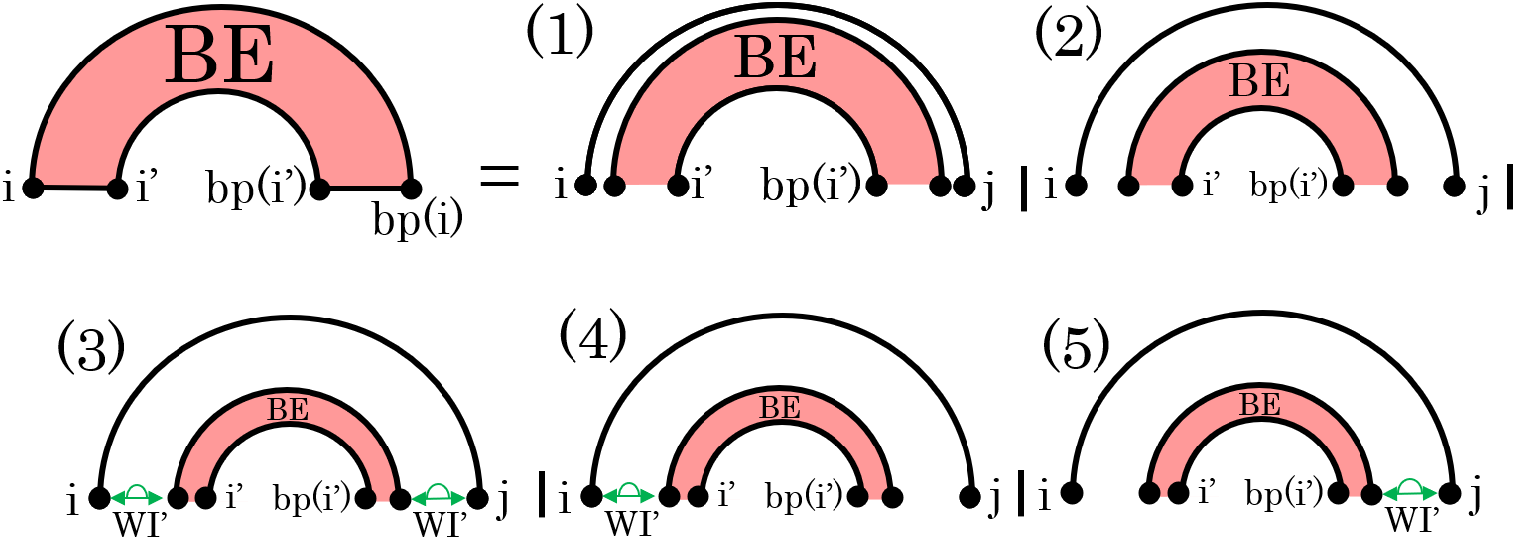
Cases of *BE*. (1) stacked pair in *G*; (2) internal loop; (3) initial and terminal weakly closed non-empty regions. (4) initial weakly closed non-empty region and terminal loop. (5) initial loop and terminal weakly closed non-empty region.

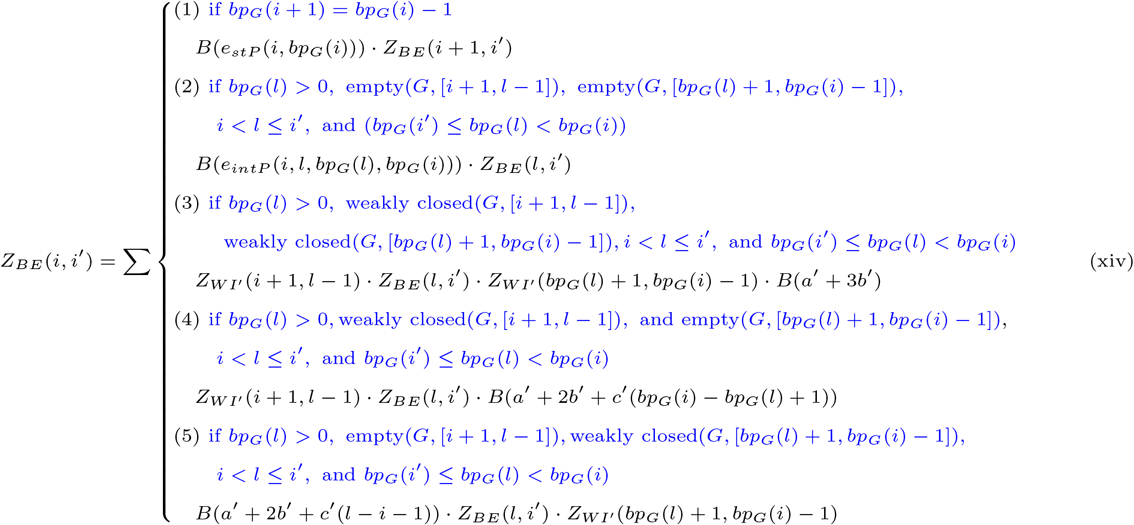

